# Spray-induced gene silencing (SIGS) as a tool for the management of Pine Pitch Canker forest disease

**DOI:** 10.1101/2024.03.05.583474

**Authors:** I.T. Bocos-Asenjo, H. Amin, S. Mosquera, S. Díez-Hermano, M. Ginésy, J.J. Diez, J. Niño-Sánchez

## Abstract

Global change is exacerbating the prevalence of plant diseases caused by pathogenic fungi in forests worldwide. The conventional use of chemical fungicides, which is commonplace in agricultural settings, is not sanctioned for application in forest ecosystems, so novel control strategies are imperative. The promising approach SIGS (Spray-Induced Gene Silencing) involves the external application of specific double-stranded RNA (dsRNA), which can modulate the expression of target genes through environmental RNA interference in eukaryotes. SIGS exhibited notable success in reducing virulence when deployed against some crop fungal pathogens, such as *Fusarium graminearum*, *Botrytis cinerea* and *Sclerotinia sclerotiorum*, among others. However, there is a conspicuous dearth of studies evaluating the applicability of SIGS for managing forest pathogens. This research aimed to determine whether SIGS could be used to control *Fusarium circinatum*, a widely impactful forest pathogen that causes Pine Pitch Canker disease. To achieve this, we designed and produced though a bacterial synthesis, dsRNA molecules to target fungal essential genes involved to vesicle trafficking (*Vps51*, *DCTN1*, and *SAC1*), signal transduction (*Pp2a*, *Sit4*, *Ppg1*, and *Tap42*), and cell wall biogenesis (*Chs1*, *Chs2*, *Chs3b*, *Gls1*) metabolic pathways. We confirmed that *F. circinatum* is able to uptake externally applied dsRNA, triggering an inhibition of the pathogen’s virulence. Furthermore, this study pioneers the demonstration that recurrent applications of dsRNAs in SIGS are more effective in protecting plants than single applications. Therefore, SIGS emerges as an effective and sustainable approach for managing plant pathogens, showcasing its efficacy in controlling a globally significant forest pathogen subject to quarantine measures.

## Introduction

In forest ecosystems, pests and pathogens play a vital role in regulating diversity. However, globalization and climate change have exposed forests to new threats, increasing reports of emerging diseases (Simler-Williamson et al., 2019). Emerging diseases are increasingly recognized by the scientific community as a significant threat to global forest ecosystems (Fisher et al., 2016; Santini and Battisti, 2019; Thakur et al., 2019). One of the greatest threats to forest health is pathogenic fungi, responsible for most emerging diseases in forest plantations (Doehlemann et al., 2017; Gomdola et al., 2022; Jayawardena et al., 2021).

It is extremely challenging to control forest tree pathogens. Unlike agricultural crops, natural forests, or forest plantations last for decades and thus, effective control strategies are required for the management of forest pathogens (Prospero et al., 2021). Currently, cultural practices or resistant species are used for disease management in forests (Prospero and Cleary, 2017; Woodcock et al., 2018). In forests, the use of chemical fungicides is subject to strict regulations and guidelines in many countries to prevent unacceptable risks to human, animal, or environmental health (Kasanen et al., 2022; Okorski et al., 2015). Repetitive applications of chemical agents in forests could also lead to resistant genotypes of pathogenic organisms or affect other non-target beneficial organisms. These reasons have prompted authorities in many countries to limit the availability of chemical agents, restrict their use only to specific pathogens in nurseries, or even ban the use of chemical fungicides in forests (Prospero et al., 2021). Consequently, the shortage of current management practices has urged the development of alternative and innovative techniques for successfully managing forest pathogens. RNA interference (RNAi)-based technologies are one of the novel and potential approaches for the control of forest pathogens and pests (Bocos-Asenjo et al., 2022; Lu et al., 2023).

RNAi is an emerging control strategy that shows great potential to manage plant diseases by silencing essential genes of pathogens. RNAi is a conserved post-transcriptional gene silencing (PTGS) mechanism that is triggered by double-stranded RNA (dsRNA) (Fire et al., 1998). It is a naturally occurring immune response and biological process in eukaryotic cells which involves the processing of double-stranded RNAs (dsRNAs) into 21-24 nucleotide (nt) small interfering RNAs (siRNAs), which then serve as guide to degrade specific messenger RNAs (mRNAs), thus suppressing the gene expression (Koch and Wassenegger, 2021; Timmons and Fire, 1998).The expression of specific genes can also be downregulated through RNAi in response to environmentally encountered dsRNA, known as environmental RNAi (eRNAi) (Tabara et al., 1998; Timmons and Fire, 1998; Whangbo and Hunter, 2008).Thus, by carefully designing dsRNAs to target genes that are essential for pathogens, eRNAi can be used to control plant diseases (Forster and Shuai, 2020; Höfle et al., 2020; Nerva et al., 2020; Werner et al., 2020). Spray-induced gene silencing (SIGS) is a novel technology based on the exogenous application of specific dsRNA molecules onto plant surfaces to protect them from pathogens (Koch et al., 2016; Wang et al., 2016). As a pathogen-specific and environmentally friendly technology, SIGS offers excellent potential for plant protection since it does not require transgenic plants, in contrast to Host-Induced Gene Silencing (HIGS) (Wang and Jin, 2017). For SIGS technology to be successful, several essential requirements must be met, including correct selection of target genes, proper design of the dsRNAs (Joga et al., 2021), as well as ensuring that the pathogens can internalize dsRNA molecules (Qiao et al., 2021).

SIGS has been extensively used against various crop fungal pathogens, such as *Botrytis cinerea* (Wang et al., 2016), *Fusarium graminearum* (Koch et al., 2016), *Sclerotinia sclerotiorum* (McLoughlin et al., 2018), *Rhizoctonia solani* (Qiao et al., 2021), *Aspergillus niger* (Qiao et al., 2021) and *Verticillium dahliae* (Song et al., 2018), or even oomycetes, such as *Phytophthora infestans* (Kalyandurg et al., 2021). In particular, research has demonstrated the use of SIGS to control fusarium head blight disease (FHB) caused by *F. graminearum* (Koch et al., 2016) and to a lesser extent by other *Fusarium* species such as *Fusarium culmorum* (Tretiakova et al., 2022), and *Fusarium oxysporum* f. sp. *lycopersici* (Ouyang et al., 2023). The significant advances of SIGS against diseases caused by agricultural fungal pathogens have urged us to explore the potential of SIGS in forestry. We have therefore investigated and tested this tool for the first time to control forest pathogens, focusing on a globally significant pine disease, pine pitch canker (PPC).

PPC disease is caused by the invasive pathogenic fungus *Fusarium circinatum* (Nirenberg and O’Donnell, 1998), considered one of the most devastating pathogens of pines worldwide (Coutinho et al., 2007; Jacobs et al., 2009; Pérez-Sierra et al., 2007), causing damping-off, shoot die-back and mortality of seedlings in nurseries, cankering, dieback and ultimately mortality in mature trees (Martín-García et al., 2017; Martínez-Álvarez et al., 2014; Viljoen et al., 1995). Globally, PPC has detrimental effects on the yield and production of timber, resulting in substantial ecological and economic losses (Wingfield et al., 2008). *F. circinatum* has been reported to infect 106 plant species, including 67 *Pinus* spp., 18 *Pinus* hybrids, 6 non-pine tree species, and 17 grass species (Carter and Gordon, 2020; Drenkhan et al., 2020; Hernandez-Escribano et al., 2018; Herron et al., 2015; Swett and Gordon, 2015; Swett et al., 2014). In addition to causing severe damage to plantations and natural forests, *Fusarium circinatum* is a significant threat to nurseries. It causes widespread losses of seedlings in forest nurseries, and as it can remain latent without developing symptoms (Martín-Rodrigues et al., 2015), it is an important source of dispersal, as infected seedlings can be planted in forests and subsequently develop the disease (Drenkhan et al., 2020). Current PPC management practices consist of phytosanitary regulations, removal of infested trees and biological control; however, they have not been sufficient to prevent biological invasion, establishment and spread of the pathogen (Eschen et al., 2015; Klapwijk et al., 2016; Liebhold et al., 2012; Martín-García et al., 2019).The harmful effects of chemical pesticides on the environment and biodiversity have motivated the search for environmentally friendly controls as the primary approach to reducing the severity of PPC disease (Iturritxa et al., 2017; Martín-García et al., 2019; Panzavolta et al., 2021). Therefore, we decided to apply SIGS technology to the protection of pine trees against PPC disease, and we have used pine seedlings to test this method since it is in nurseries where the dissemination and incidence of *F. circinatum* begins.

By designing and applying external dsRNAs targeting essential genes of the pathogen, we demonstrated that *F. circinatum* infection in pine seedlings can be successfully controlled. This is the first time SIGS has been used to control fungal disease in trees. Promising results suggest that implementing SIGS technology might be applicable to control many other forest diseases.

## Materials and Methods

### Target gene selection and dsRNA design

Eleven fungal genes were selected as RNAi targets based on their use in previous RNAi studies and their relevance for fungal virulence, pathogenicity, and survival (table 1). Three of these genes, the vacuolar protein sorting 51 (*Vps51*), the large subunit of the dynactin complex (*DCTN1*), and a suppressor of actin (*SAC1*) –like phosphoinositide phosphatase, are involved in the vesicle trafficking pathway (Cai et al., 2018). Four of them, three 2A phosphatases (*Pp2a*, *Sit4, Ppg1*) and the phosphatase 2A-associating protein (*Tap42*) belong to a signal transduction pathway (Yu et al., 2014); and the remaining four, the chitin synthases (C*hs1,* C*hs2, Chs3b*) and the β-1,3-Glucan Synthase (*Gls1*), participate in cell wall biogenesis (Chen et al., 2016a; Cheng et al., 2015). Orthologous *F. circinatum* genes were identified using the protein sequences of selected genes and the *F. circinatum* NCBI-indexed genome (ASM2404739v1) in the online NCBI blastptool.

**Table 1.** Target genes in *Fusarium circinatum*.

| Pathway | Gene | Encoded protein | Protein ID | Reference |
| --- | --- | --- | --- | --- |
| Vesicle trafficking | *FcVps51* | Hypothetical protein FCIRC_4363 | KAF5683607.1 | (Cai et al., 2018) |
|  | *FcDCTN1* | Tip elongation 1 | KAF5656821.1 | (Cai et al., 2018) |
|  | *FcSAC1* | Phosphatidylinositide phosphatase SAC2 | KAF5666971.1 | (Cai et al., 2018) |
| Signaling | *FcPp2a* | Serine threonine phosphatase PP2A catalytic subunit | KAF5688398.1 | (Yu et al., 2014) |
|  | *FcPpg1* | Serine/threonine-protein phosphatase PP2A catalytic subunit | KAF5683879.1 | (Yu et al., 2014) |
|  | *FcSit4* | Serine/threonine-protein Phosphatase PP1-1 | KAF5690545.1 | (Yu et al., 2014) |
|  | *FcTap42* | TAP42 component of the Tor signaling pathway | KAF5688382.1 | (Yu et al., 2014) |
| Cell wall biogenesis | *FcChs1* | Chitin synthase 1 | KAF5654678.1 | (Cheng et al., 2015) |
|  | *FcChs2* | Chitin synthase 2 | KAF5678417.1 | (Cheng et al., 2015) |
|  | *FcChs3* | Chitin synthase 3 | KAF5658513.1 | (Cheng et al., 2015) |
|  | *FcGls1* | 1,3-beta-D-glucan synthase subunit | KAF5676366.1 | (Chen et al., 2016a) |

For dsRNAs design, stretches between 200 and 250 nucleotides (nt) of the *F. circinatum* genes were selected by assessing RNA target site accessibility using 8 and 16 nt with the NAplfold v2.6.4 (Bernhart et al., 2011; Lorenz et al., 2016), available through the RNAxs Webserver (default parameters). The output was fed into a custom-made R script to obtain plots of site accessibility throughout the complete mRNA target sequence. Site accessibility was scored by a penalty according to a pre-determined accessibility threshold (0.011 for 8-nt and 0.001 for 16-nt). Regions with the highest score were then chosen by visual inspection of the plots. To prevent off-target effects on beneficial microbes or plants, we avoided using DNA stretches falling within conserved gene regions. The dsRNAs were aligned to the host transcriptome using the online NCBI nblast tool to ensure no off-target effects in the host. Three DNA constructs containing DNA stretches in each pathway and restriction *kpn*I (GGTACC), *Bgl*II (AGATCT), or *Sac*II (CCGCGG) sites at the constructs 3’– and 5’-ends were designed. These constructs corresponded to dsRNAs: [1] dsRNA-*VDS,* targeting *Vps51, DCTN1, and SAC1 genes*; [2] dsRNA-*PTP,* targeting *Pp2a*, *Sit4, Ppg1,* and *Tap42*; and [3] dsRNA-*CHS,* targeting C*hs1,* C*hs2, Chs3b,* and *Gls1*. A DNA construct corresponding to a dsRNA targeting the Yellow Fluorescent Protein (*YFP*) gene was also included as a negative control for the dsRNA bioassay (dsRNA-*YFP*). Finally, resulting dsRNA molecules had lengths of 517 nt (dsRNA-*YFP*), 498 nt (dsRNA-*VDS*), and 800 nt (dsRNA-*PTP* and dsRNA-*CHS*), including fragments of 165 to 200 nt of each gene (table S1 “dsRNA sequence”).

### dsRNA production using a genetically modified bacteria

For dsRNA vectors assembly, constructs sequences were synthesized in Integrated DNA Technologies (IDT, Coralville, IA, USA) and cloned into the T777T plasmid (Addgene plasmid # 113082; http://n2t.net/addgene:113082; RRID:Addgene_113082) which contains two IPTG-inducible T7 promoter (5’-TAATACGACTCACTATAG-3’) in opposite directions that synthesize complementary RNA strands to form dsRNAs (Sturm et al., 2018).Plasmid and inserts were digested with *Kpn*I and BglII or *Sac*II restriction enzymes (New England Biolabs, Ipswich, MA, USA) following manufacturer instructions. Then, they were ligated in 20 µL reactions with the enzyme T4 DNA ligase (Thermo Fisher, Waltham, MA, USA) to get the pT777T-*FcVDS,* pT777T-*FcPTP,* pT777T-*FcCHS* and pT777T-*YFP* vectors (table S1 “Plasmid map”).

Plasmids were cloned into TOP10 Chemically Competent *Escherichia coli* cells by heat shock method (42°C, 50 s) for plasmid propagation. Transformants carrying recombinant plasmids were screened in Luria-Bertani (LB) plates [containing 10 g Tryptone (Scharlab S.L., Barcelona, Spain), 10 g NaCl (Panreac Química S.L.U, Spain), 5 g Yeast Extract (Scharlab S.L., Barcelona, Spain), 15 g Agar-Agar (Panreac Química S.L.U, Spain) per L] supplemented with ampicillin at 100 µg/L (Amp_100_). Plasmids were purified using the NucleoSpin Plasmid kit (Macherey-Nagel^TM^) and cloned, as before, into the RNaseIII-null mutant *E. coli* strain HT115(DE3) able to produce dsRNA (Dasgupta et al., 1998; Timmons et al., 2001) which has been proven to be biological active against plant pathogenic fungi (Niño-Sanchez et al., 2021). Transformants (HT115/*FcVDS*, HT115/*FcPTP*, HT115/*FcCHS,* and HT115/*YFP*) were screened as described and grown in 15 mL flasks containing 5 mL of LB liquid medium with Amp_100_ at 37°C at 270 rpm to a 600 nm optical density (OD_600_) of 0.8. Cultures were supplemented with isopropyl-β-D-1-thiogalactopyranoside (IPTG) at a final concentration of 1.0 mM to induce dsRNA production and incubated under the same conditions until reaching the stationary phase (between 4 and 8 h). Bacterial cells were concentrated by centrifugation at 4400 x g for 5 min) and used for total RNA extraction with TRIzol® (Thermo Fisher, Waltham, MA, USA) using ethanol washes at 65% ethanol instead of 75% to enrich the dsRNAs. Genomic DNA and single-stranded RNA were removed from the dsRNA preparations by digestion for 45 min at 37°C with 1 µL DNase I Ambion ^TM^ (2 U/µl) (Thermo Fisher, Waltham, MA, USA) and 1µL RNase T1 (1000 U/µl) (Thermo Fisher, Waltham, MA, USA) in x1 DNase buffer (Thermo Fisher, Waltham, MA, USA). Enzymes were removed using the RNA Clean & Concentrator Kit (Zymo Research, Irvine, CA, USA), and dsRNA production was verified in agarose gels at 1 % run at 100 V for 40 min.

### Fungal isolate and plant material

*F. circinatum* isolate Fc072v used here was isolated in a previous study from a symptomatic *Pinus radiata* located in Cantabria (Northern Spain) (Zamora-Ballesteros et al., 2021). For spore production, Fc072v was grown in a 250 mL flask containing 100 mL of GOX liquid medium [60 g of sucrose (Sigma Aldrich, St. Louis, MO, USA), 7 g of NaNO_3_ (Sigma Aldrich, St. Louis, MO, USA), 3 g of peptone (Condalab, Madrid, Spain), 1 g of KH_2_PO_4_ (Panreac Química S.L.U., Barcelona, Spain), 0.5 g of MgSO_4_·7H_2_O (Panreac Química S.A, Spain), and 0.5 g of KCl (Panreac Química S.L.U, Barcelona, Spain), per L at a pH of 7] for 48 h at 180 rpm and 25°C. Cultures were filtered twice through sterile cheesecloth for hyphal removal, and spores were collected by centrifugation at 4400 x g for 10 min. *P. radiata* seedlings were grown from seeds (Provenance: Galicia, Spain) in a growth chamber at 21.5°C, with a 16/8 h light/dark photoperiod. Baby spinach (*Spinacia oleracea*) detached leaves, used in the rapid efficacy test, were obtained fresh from a local market.

### Fungal uptake evaluation

To assess the uptake ability of the pathogen, we followed the methodology described in Hamby et al. (2020). Briefly, Fluorescein-labeled dsRNA was synthesized using 2 µL of the plasmid pT777T-*YFP* in 20 µL reactions of the MEGAscript^TM^ RNAi Kit (Thermo Fisher, Waltham, MA, USA) with 2 µL of the Fluorescein labeling mix (Roche, Basel, Switzerland), instead of the recommended 2 µL of each NTP. *F. circinatum* spore suspension was seeded in 5 mL of Potato Dextrose Broth (PDB) (Scharlab S.L., Barcelona, Spain) at a final concentration of 5 × 10^5^ spores/mL and incubated for 8 h at 80 rpm and 23°C. Then, 5 µL of the germinated spores were placed on the surface of microscopy slides containing 3 mL of Potato Dextrose Agar (PDA) (Scharlab S.L., Barcelona, Spain) and treated with 10 µL of fluorescein-labelled dsRNA at a concentration of 100 ng/µL. The Slide cultures were kept in the dark for 3 h in an opaque box on moistened filter paper to maintain humidity. Slides were treated with 20 U of Micrococcal nuclease enzyme (MNase; Thermo Fisher, Waltham, MA, USA) for 30 min at 37°C to remove external fluorescein-labelled dsRNAs (Qiao et al., 2021). Then, they were analyzed in a laser scanning confocal microscopy (LSCM) (Leica SP8) at an excitation of 491 nm and detection of 500-550 nm.

### Rapid dsRNAs-efficacy assay in spinach leaves

A methodology for a rapid assessment of the designed dsRNAs was developed using detached spinach leaves. This method was intended as a preliminary evaluation to assess the designed dsRNA molecules before their analysis in the *F. circinatum* primary host (*P. radiata*). For this assay, 10 µL drops of the dsRNA treatments (dsRNA-*VDS*, dsRNA-*PTP*, dsRNA-*CHS*) or dsRNA-*YFP* control suspensions at 100 ng/ µL were applied to the abaxial left or right sides of the detached spinach leaf’s midrib. Then, leaves were inoculated on the same side as the dsRNA treatments and dsRNA-*YFP* control with 10 µL of 10^5^ *F. circinatum* spore/mL suspension and incubated on moist paper at room temperature in a tightly sealed container at 25°C. Three independent evaluations, each including 30 leaves per treatment, were carried out for the lesion size evaluations. Each leaf was considered as a biological replicate. Necrotic lesion diameters were measured at 4 days post-inoculation (dpi). The leaves which could not be measured for technical reasons were withdrawn from the experiment before the statistical analysis. For fungal biomass evaluations, four leaves were pooled into compound samples, each compound sample constitutes a biological replicate. Two compound samples were taken from each experiment for a total of six biological replicates.

### dsRNAs-efficacy against Fusarium circinatum in Pinus radiata

For assessing the dsRNA control against *F. circinatum* in pine seedlings, a wound was made 1 – 1.5 cm above the collar of *P. radiata* seedlings following the methodology of Zamora-Ballesteros et al. (2021). Wounds were inoculated with 5 µL of a suspension of 10^5^ *F. circinatum* spores/mL and treated with 5 µL of 300 ng/µL suspensions of the dsRNA treatments or the dsRNA-*YFP* control. The aerial part of each seedling was also treated by spraying 1 mL of 300 ng/µL suspensions of the dsRNA treatments or the dsRNA-*YFP* control. Inoculated seedlings were kept in a phytotron at 21.5°C, with a 16/8 h light/dark photoperiod for 35 days. Symptom progression was assessed visually at 25 and 35 dpi using a modified version of a 1 to 5 disease score scale (Correll et al., 1991) (figure 1). In this scale, 0 represents a healthy plant with no symptoms; 1 the appearance of resin or necrosis at the point of inoculation and healthy foliage, 2 the appearance of resin or necrosis at the point of inoculation, green foliage and slight damping-off; 3 noticeable damping-off symptoms, slight wilting and yellowing in the tip; 4 severe wilting and yellowing and noticeable dieback; and 5 a dead plant. This scale was modified with intermediate values to improve the precision of the disease score measurement and to increase its accuracy. The experiment was carried out using 35 four-month-old *P. radiata* seedlings per treatment, with each seedling representing a biological replicate. Mock-inoculated (H_2_O) seedlings served as a negative control to compare symptom development. Two positive controls were used: plants inoculated only with *F. circinatum* without dsRNA treatment and plants inoculated with *F. circinatum* and treated with the non-pathogen-specific dsRNA control (dsRNA-*YFP*). The latter was included to check whether the effect of the dsRNA treatments is related to the sequence specificity of the designed molecules. dsRNA-*VDS*, dsRNA-*PTP*, dsRNA-*CHS* were tested alone and in combination.

**Figure 1.**
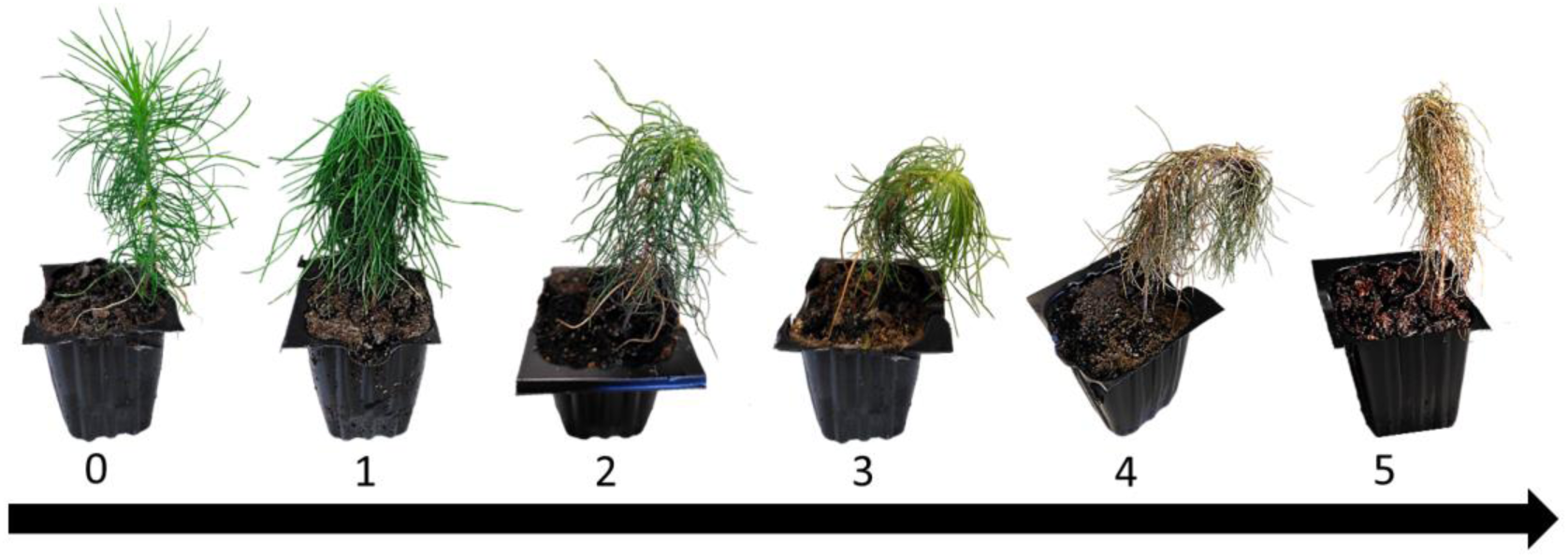
The seedlings were visually scored for disease symptoms according to a 0–5 disease score empirical scale modified from Correll et al. (1991) to increase its accuracy: 0 = healthy plant or no symptoms; 1 = resin and/or necrosis at the point of inoculation and healthy foliage; 2 = resin and/or necrosis beyond the point of inoculation, green foliage and slight dumping-off; 3 = noticeable damping-off symptoms, slight wilting and yellowing in the tip; 4 = severe wilting and yellowing and noticeable dieback; 5 = dead plant. Intermediate points have also been taken to provide more extensive information and accuracy of measurements.

To assess whether periodic application of dsRNA treatments could improve SIGS efficacy, reduce or even control disease progression, a second trial was conducted in which seedlings were treated with dsRNAs at 0 dpi (single application), and then half of them were also treated every 7 days up to 30 dpi (weekly). This experiment exclusively included the dsRNA-*CHS* treatment, which was selected based on the first trial, and the dsRNA-*YFP* and mock-inoculated (H_2_O) controls. Three independent experiments were carried out, each using 10 seedlings per treatment or control, inoculated and incubated as before. Symptoms were evaluated 30 dpi according to the modified (Correll et al., 1991) scale.

### Fungal biomass quantification

The fungal biomass was estimated with quantitative PCR (qPCR) for assessing fungal proliferation. For this purpose, plant material was collected at 4 dpi from the spinach and pines assays (Section 2.5 and 2.6). Total genomic DNA was extracted using a CTAB protocol (Chen and Ronald, 1999).1 µL of the extracted DNA was used in 20 µL reactions of Phusion™ High-Fidelity DNA Polymerase (Thermo Fisher, Waltham, MA, USA) containing 0.4 µL of 1:1000 SYBR™ Green I Nucleic Acid Gel Stain (Thermo Fisher, Waltham, MA, USA). Primers *FcRPB2*-q-F and *FcRPB2*-q-R were used to amplify the *F. circinatum* RNA polymerase II gene RPBII (*RPB2*) at 1 µM (table S2). Similar reaction with the primers *SoActin*-q-F and *SoActin*-q-R weas performed to amplify the spinach *Actin* gene (table S2). Amplifications were carried in the QuantStudio 6 Flex Real-time PCR System (Applied Biosystems) using a protocol consisting of an initial activation step at 94°C for 2 min 30 sec, and 40 cycles of 94°C for 15 s, 60°C for 20 s, and 72°C for 30 s. Ct values were determined with the QuantStudio Real-Time PCR v1.7.1 software. We correlated the amplification levels of the fungal *RPB2* gene with those of the *actin* spinach gene to calculate the relative amount of fungal growth. Gene relative quantification was determined via the 2-ΔΔCt method by normalizing the amount of target transcript to the number of spinach– or pine-reference genes (Livak and Schmittgen, 2001). Cts from plant material treated with the dsRNA– *YFP* control was used as reference against the dsRNA-treated samples (Wang et al., 2016).

### RNA-seq assay and differential expression analysis

For assessing the effect of the dsRNA-*PTP* (which presumable affects the modulation of transcriptional responses) on the *F. circinatum* transcriptome, four 4-month-old dsRNA-treated or mock-inoculated (H_2_O) *P. radiata* seedlings were inoculated with 10^5^ spores/mL of *F. circinatum* strain Fc072v. We only used dsRNA-*PTP* for transcriptomic analysis, as genes in signal transduction pathways can also regulate other essential pathways (Yu et al., 2014). Spore inoculation, dsRNA treatment and pine seedlings incubation were carried out as before (section 2.6). At 4 dpi, three samples consisting of both treated and mock seedlings were collected and used for total RNA extraction with the Sigma mirPremier® microRNA Isolation Kit (Sigma Aldrich, St. Louis, MO, USA) following the manufacturer’s instructions. RNA was sent to Macrogen Incorporation (South Korea) for paired-end sequencing using an Illumina NextSeq platform. Library preparation was performed by an external provider (Macrogen Inc., South Korea) with the TruSeq Stranded mRNA library kit. The raw data was assessed for quality control using FastQC v0.11.9 (Andrews, 2010). Then, it was used for trimming Illumina adaptor sequences, and removing low-quality reads, contaminant DNA, and PCR duplicates with Trimmomatic v0.38 (Bolger et al., 2014). Trimmed reads were mapped to the transcriptome of *F. circinatum* strain CMWF1803 (Assembly ASM2404739v1, NCBI Acc. Number GCA_024047395.1) using the splice-aware aligner HISAT2 v2.1.0 (Kim et al., 2015). After assembly, expression profiles (abundance of genes/transcripts) were represented as read counts and normalization values, which were calculated based on transcript length and depth of coverage. Normalization values were provided as FPKM (Fragments Per Kilobase of transcript per Million mapped reads). The resulting count files were formatted into a count matrix suitable for differential expression (DE) analysis. Raw reads have been deposited in the NCBI SRA Database under accession numbers SAMN38792722 to 29 (BioProject PRJNA1051661).

For the DE analysis, the counts’ matrix tables were loaded into the software R v4.2.1 (R Core Team, 2022), and analyzed using the DESeq function of the package DESeq2 v1.34.0 (Love et al., 2014). Pairwise comparisons for RNA treatment vs control were performed to identify differentially expressed genes using a threshold of log2(|Fold-change|) ≥ 1 at a false discovery rate (FDR) < 0.05.

### Statistics

Differences in disease score in pine assays were analyzed using ordered logistic regression (OLR). Variables considered in this study were treatment type (negative and positive controls and dsRNA solutions), assessment days post-inoculation (4 dpi, 25 dpi or 35 dpi) and frequency of treatment application (single or weekly). The assumption of proportional odds was checked using the Brant test (not significant, *p*-val ≥ 0.05 for all variables). Differences in lesion size and fungal biomass between dsRNA-treated and non-treated spinach leaves were assessed using the Wilcoxon rank test. Statistical significance was considered for *p*-values < 0.05.

## Results

### *Fusarium circinatum* is able to uptake externally applied dsRNA

To determine whether the dsRNA is taken up and internalized by the fungal structures of the pathogen *F. circinatum*, it was incubated with fluorescein-labeled dsRNA and then visualized by Confocal Laser Scannig Microsopy (CLSM). The presence of fluorescent signals inside germinating spores of *F. circinatum* 3 h after incubation with fluorescein-labeled dsRNA confirmed the incorporation of these molecules into the fungal structures (figure 2). Furthermore, fluorescence was observed across multiple Z-stacks. Thus, dsRNA was located inside of the mycelium in different focal planes within germinating spores, which indicated that the dsRNA was internalized right after spore germination (figure S1). Therefore, our results demonstrated that *F. circinatum* successfully uptakes externally applied dsRNAs and suggest it is susceptible to SIGS.

**Figure 2.**
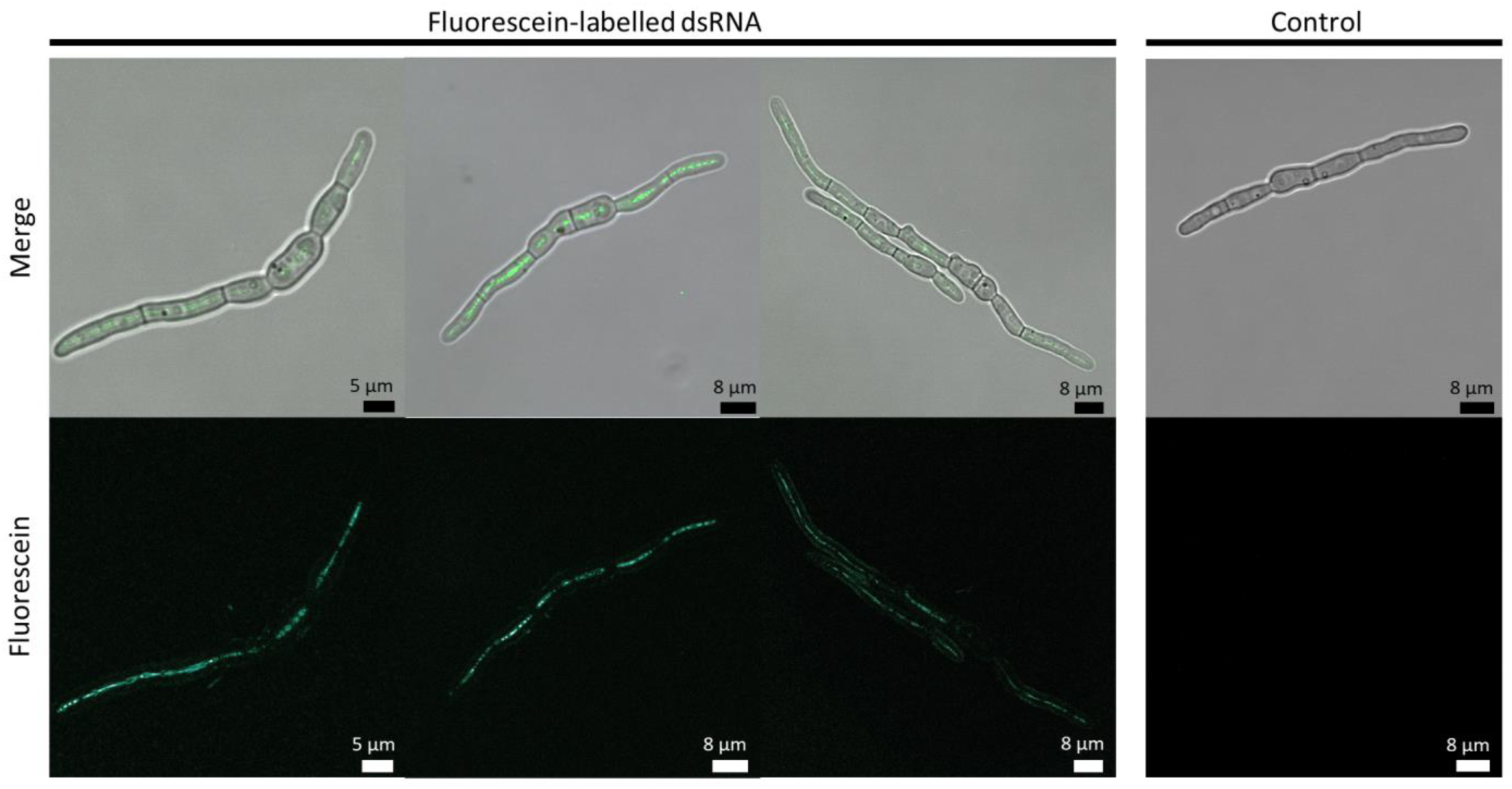
*Fusarium circinatum* is able to take up externally applied dsRNA*. F. circinatum* germlings were treated with fluorescein-labelled dsRNA 3 hours before visualization by means of CLSM. MNase was used to degrade external labelled dsRNA and ensure the absence of dsRNA adhesion to the fungal cell wall. Absence of fluorescent signal is shown in untreated *F. circinatum* germlings (images on the right).

### Identification and validation of *Fusarium circinatum* target genes for its use in SIGS technology

dsRNAs in this study were specifically designed to target up to four genes in metabolic pathways critical for the survival of the fungus. Three metabolic pathways were finally chosen for the design of the dsRNA molecules: (1) vesicle trafficking, (2) the signal transduction pathway (cell signaling), and (3) cell wall biogenesis. Essential genes in these pathways were identified based on their successful use in previous RNAi studies (section 2.1). Based on this selection, we constructed three dsRNA molecules (dsRNA-*VDS*, dsRNA-*PTP*, dsRNA-*CHS*), and a molecule with an unspecific target to be used as a control (dsRNA-*YFP*). Then, we conducted in-plant trials to evaluate the efficacy of these dsRNA molecules in reducing the virulence of *F. circinatum* by SIGS. Considering the time requirements associated with working with pines, we needed a quick methodology to test the efficacy of the dsRNA molecules. To achieve this, we used detached spinach leaves since they are susceptible to *F*. *circinatum* infection, and symptoms are easy to assess. As a result, we were able to quickly determine whether the designed molecules protected against the pathogen.

Treatments with dsRNA-*VDS*, dsRNA-*PTP*, and dsRNA-*CHS* had a protective effect on detached spinach leaves (figure 3). The development of necrotic lesions was reduced in all dsRNA treatments compared to the dsRNA-*YFP* treated control (figure 3A). Disease severity (based on the mean lesion size) was reduced by 43% (*p* < 0.01) in dsRNA*-VDS* treatments, 33% (*p* < 0.01) in dsRNA*-PTP* treatments, and 35% (*p* < 0.01) in dsRNA*-CHS* treatments, compared to the dsRNA-*YFP*-treated control (figure 3B). Fungal biomass quantification performed by qPCR confirmed the reduction of fungal growth on leaves treated with the designed dsRNAs: the relative amount of fungal growth was reduced in the treatments by 46% (*p =* 0.06) in dsRNA-*VDS*, 40% (*p* < 0.01) in dsRNA-*PTP* and 36 % (*p* < 0.01) in dsRNA-*CHS* compared to the dsRNA-*YFP* control (figure 3C). Therefore, the assessed dsRNAs reduced the fungal growth and thus the disease incidence on detached spinach leaves. Furthermore, it was shown that spinach leaf assays are a reliable and rapid method to test the efficacy of various dsRNAs against *F. circinatum*.

**Figure 3.**
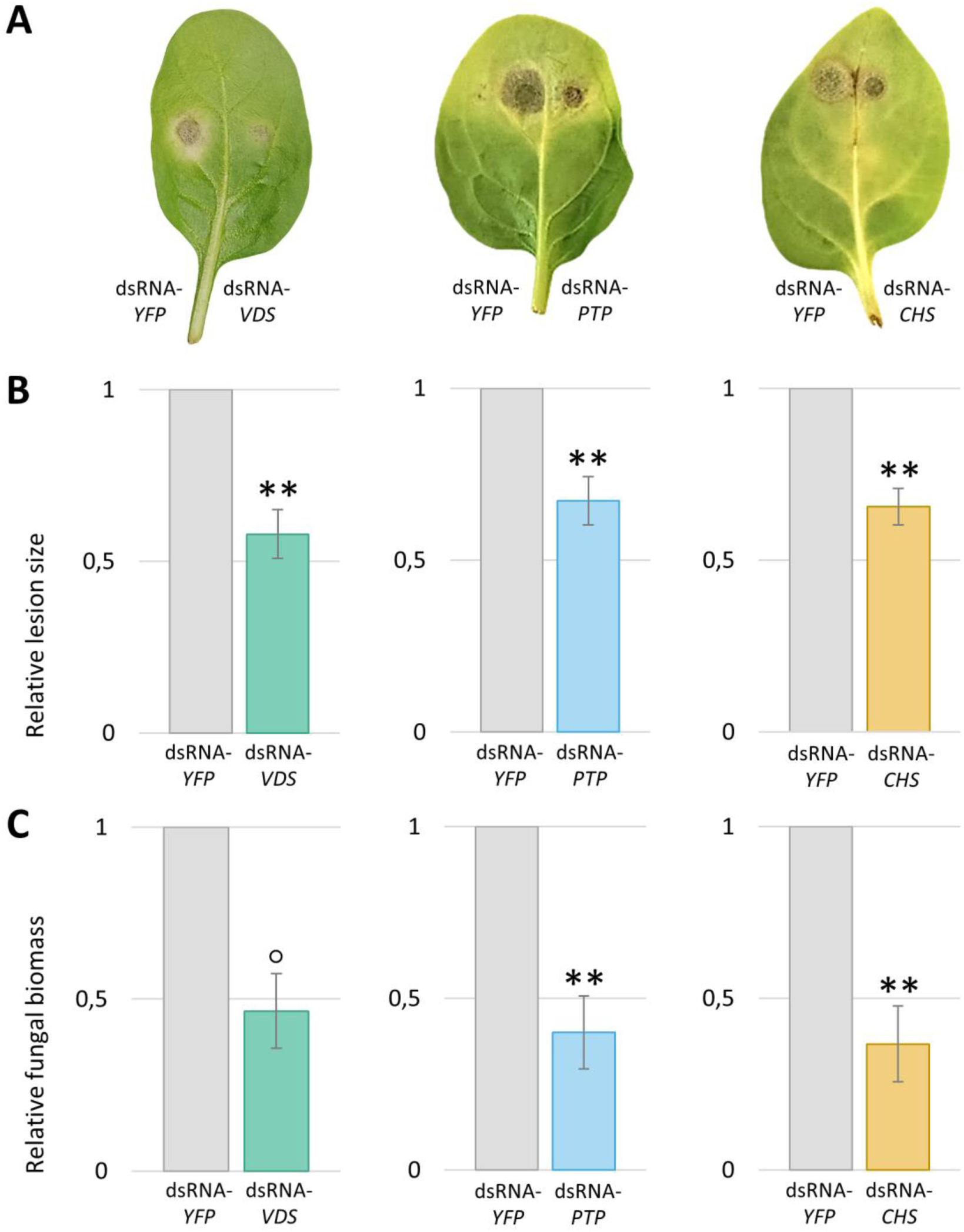
Application of designed dsRNAs against essential genes reduces the virulence of *Fusarium circinatum* in detached spinach leaves. (A) Representative detached spinach leaves show smaller necrotic lesion size in the assessed treatments (dsRNA*-VDS/*dsRNA-*PTP/*dsRNA-*CHS)* as compared to the control treatment (dsRNA-*YFP*) at 4 dpi. (B) Relative lesion size depicts the reduction of disease severity in the applied treatments (dsRNA-*VDS/*dsRNA-*PTP/*dsRNA-*CHS)* as compared to the control treatment (dsRNA-*YFP*) at 4 dpi, which were measured with the help of an electronic calibrator, assigning a value of 1.0 to the lesion size area in the control treatment. (C) Quantification of fungal biomass by means qPCR shows a reduction of *F. circinatum* in dsRNAs treatments (dsRNA-*VDS/*dsRNA-*PTP/*dsRNA-*CHS)* as compared to the control treatment (dsRNA-*YFP*). Level of statistical significances is determined by non-parametric Wilcoxon rank test. Bars and error bars represent mean relative values ± SEs between biological replicates. The level of statistical significances was determined by non-parametric Wilcoxon rank test (°: *p* < 0.10, **: *p* < 0.01)

### SIGS mediated control of *F. circinatum* in pine seedlings

To explore the efficacy of dsRNA-*VDS*, dsRNA-*PTP,* dsRNA-*CHS,* and a mixture of the three to control PPC caused by *F. circinatum* in its natural host, dsRNA treatments were tested for their ability to reduce virulence in four-month-old *P. radiata* seedlings. The fungal virulence on pine seedlings was evaluated at two different time points (25 dpi and 35 dpi) according to a disease score scale modified from Correll et al. (1991). Mock-inoculated seedlings remained healthy according to the disease scale 0 ± 0 (mean ± SD) for the entire duration of the trial. Inoculated control plants showed a mean disease score of 2.38 ± 0.48 at 25 dpi and 3.74 ± 0.66 at 35 dpi, whereas the mean disease score of inoculated plants treated with dsRNA-*YFP* control was 2.54 ± 0.57 at 25 dpi and 3.77 ± 0.61 at 35 dpi. Therefore, no differences were observed between these two positive control treatments (*p*-value 25 dpi = 0.83; *p*-value 35 dpi = 0.99). dsRNA-*YFP-*treated positive control was used as a reference for statistical calculations. At 25 dpi, the virulence of *F. circinatum* in seedlings treated with dsRNA*-VDS*, dsRNA-*PTP*, dsRNA-*CHS,* and dsRNA-mix was reduced compared to the control. The greatest reduction at this time point was observed in the dsRNA-*CHS* treatment (mean = 1.49 ± 0.77), which decreased virulence by 48% compared to the dsRNA-*YFP*-treated control (*p*-value < 0,01 according to OLR test). The dsRNA-*PTP* (mean = 1.67 ± 0.8) and dsRNA-mix (mean = 1.8 ± 0.80) treatments reduced pathogen virulence by 24% and 29% respectively (*p*-value < 0.01 according to ORL test) (figure 4, table S3). At 35 dpi all treatments showed statistically significant differences compared to the control group, with symptom reductions ranging from 14% to 23% (*p*-value < 0.01 according to OLR test) depending on the treatment and dsRNA-*CHS* (mean = 2.9 ± 0.56) being the most effective in controlling the disease again (figure 4, table S3).

**Figure 4.**
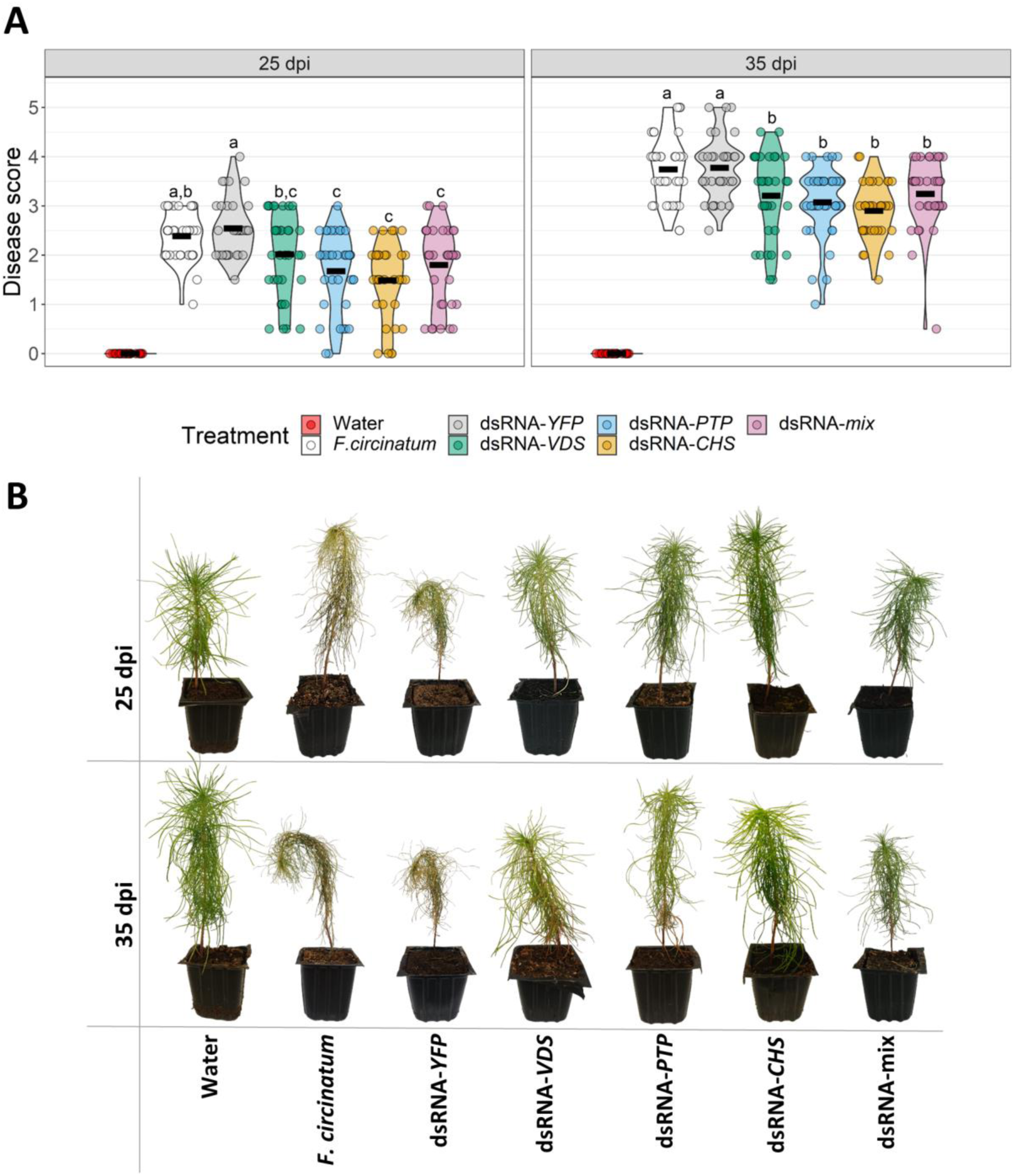
dsRNAs treatments inhibit *Fusarium circinatum* virulence in *Pinus radiata* young plants. (A) Violin plot showing the distribution of disease scores for each treatment applied according to the scale at two time points (25 dpi and 35 dpi). Black horizontal bars depict the mean disease score, and each point represents a biological replicate (n=35). Statistically significant differences are indicated by different letters determined by OLR test (*p* < 0.05). (B) Representative images taken at two time points (25 dpi and 35 dpi) of *P. radiata* plants infected with *F. circinatum* and treated.

Although there was no statistical difference between the dsRNA treatments, numerically dsRNA*-CHS* provided the greatest protection with one treatment application yielding a virulence reduction of 23% compared to the dsRNA-*YFP* control at 35 dpi. To assess whether we could increase the effect of this treatment by providing prolonged protection, we decided to increase the number of applications using a weekly application regime for 35 days. The disease score after weekly application of dsRNA-*YFP* and dsRNA-*CHS* treatments was 3.55 ± 0.80 and 2.23 ± 0.81, respectively (figure 5A). Thus, increasing the number of applications of dsRNA-*CHS* reduced pathogen virulence by 37% compared to the dsRNA-*YFP* control (*p*-value < 0.01 according to OLR test) (figure 5B, table S4). Furthermore, weekly treatment with dsRNA-*CHS* provided higher protection compared with a single dsRNA-*CHS* treatment applied once on the day of infection, with mean disease scores of 3.55 ± 0.80 and 2.83 ± 0.94, respectively (figure S2). Weekly treatment reduced *F. circinatum* virulence by 21% compared to a single application (*p*-value < 0.05 according to OLR test) suggesting that increasing the number of applications provides higher levels of protection (table S5).

**Figure 5.**
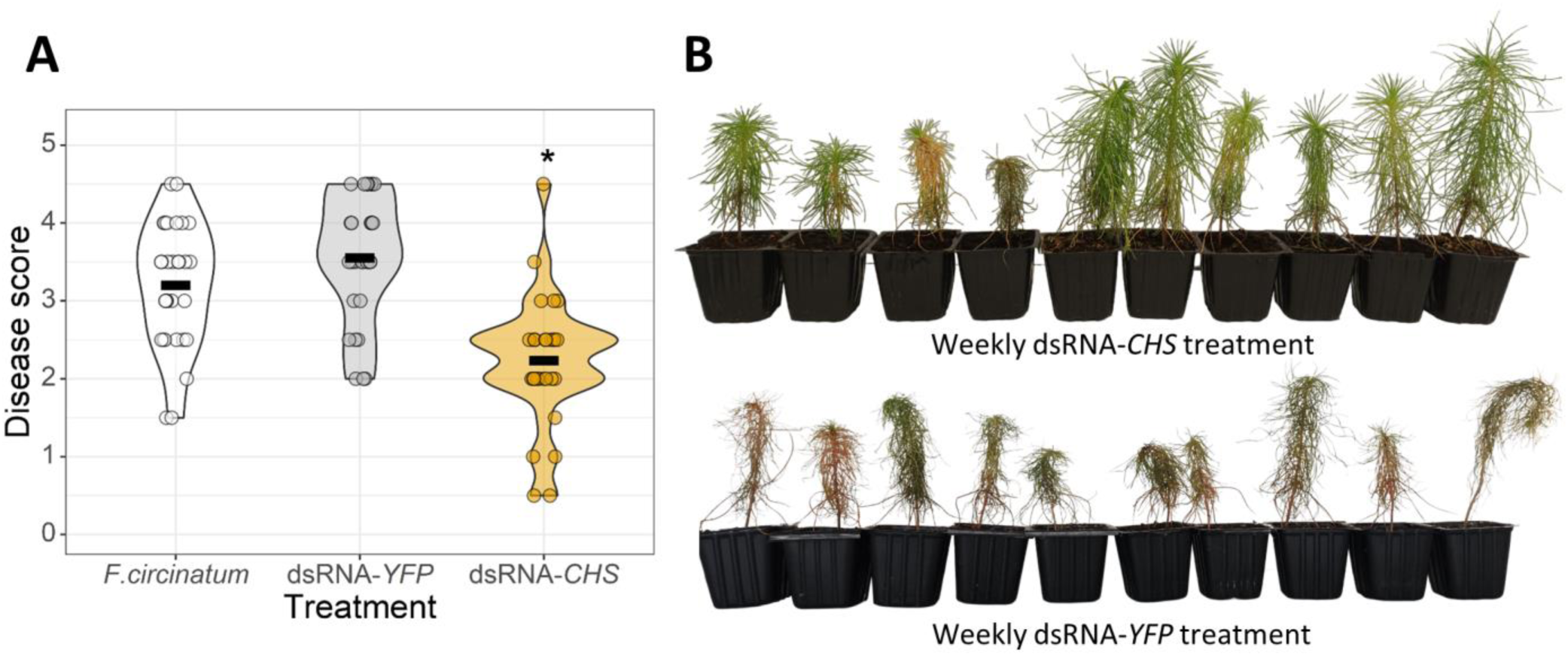
Weekly dsRNA-*CHS* application enhances the efficacy of SIGS protection against *Fusarium circinatum*. Pine seedlings were treated with dsRNA weekly and evaluated at 30 dpi according to the scale (A) Violin plots representing the distribution of disease severity data at 30 dpi of pines treated weekly with dsRNA-*CHS*. Black horizontal bars depict the mean disease score. Statistically significant differences are analyzed by OLR test and represented by asterisks (*: *p* < 0.05). (B) Pine seedlings infected with *F. circinatum* and treated weekly with dsRNA-*CHS* and dsRNA-*YFP* (control) at 35 dpi.

### Targeting genes involved in signal transduction through SIGS leads to significant transcriptomic alterations

We determined the transcriptomic alterations in treated and mock seedlings of pine at 4dpi after exogenous application of dsRNA-*PTP* through RNA-seq. Almost 21 genes were down-regulated, and 3 genes were up-regulated in treated samples and their ontology was analyzed. 11 of the down-regulated genes are putatively involved in pathogenicity and virulence of *F. circinatum*. Those genes are *Endoglucanase c* (FCIRC_4841)*, Hypothetical Protein* (*lysozyme family protein*) (FCIRC_ 8313)*, Snf5 subunit* (FCIRC_9605)*, Tripeptidyl aminopeptidase* (FCIRC_9078)*, Metalloprotease MEP1* (FCIRC_5870)*, 2,3-bisphosphoglycerate-dependent phosphoglycerate mutase* (FCIRC_1245)*, Transcriptional regulator* (FCIRC_4556)*, Aliphatic nitrilase* (FCIRC_7774) *, Major facilitator superfamily transporter (MFS)* (FCIRC_989)*, GAP1-general amino acid permease* (FCIRC_4383), and *Acetyl hydrolase* (FCIRC_9937) and are known to be involved in various biological processes (metabolic activity, transferase activity, transport, hydrolase activity and oxidoreductase) and cellular components (cell wall biogenesis or organization) (table 2).

**Table 2.**
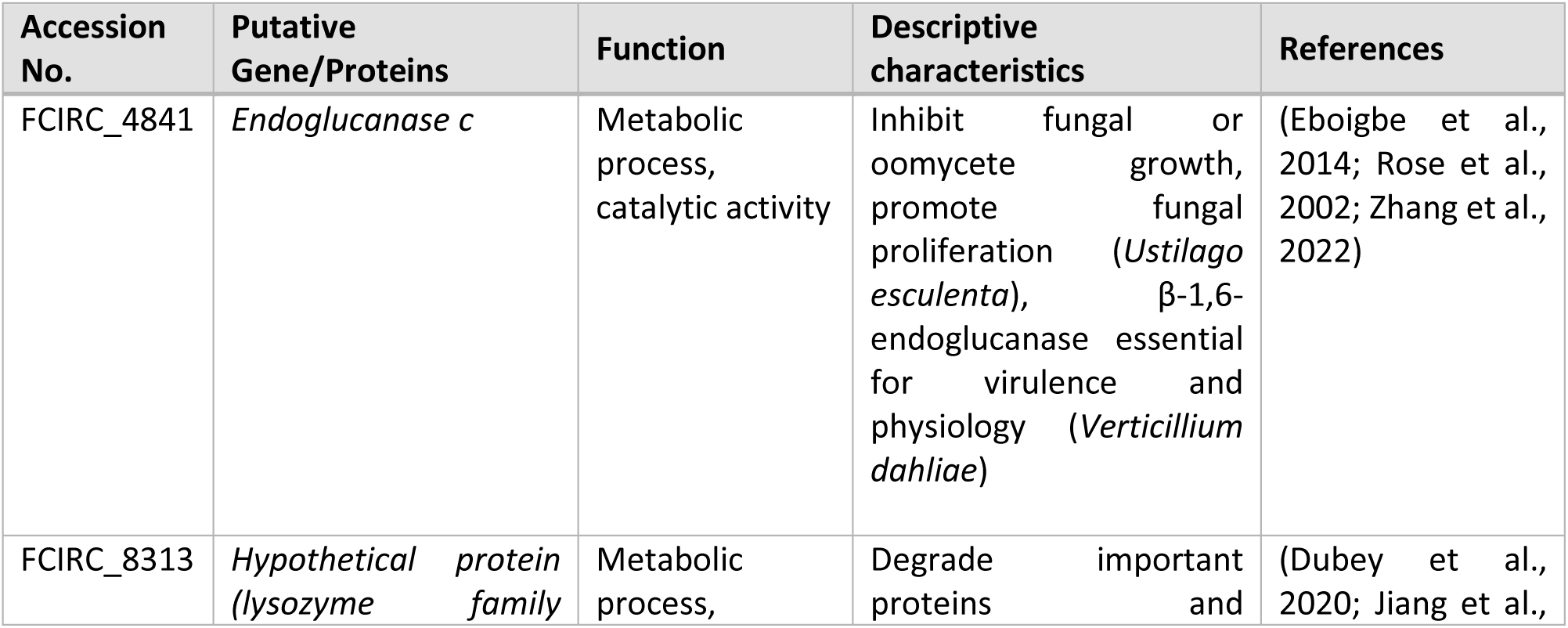
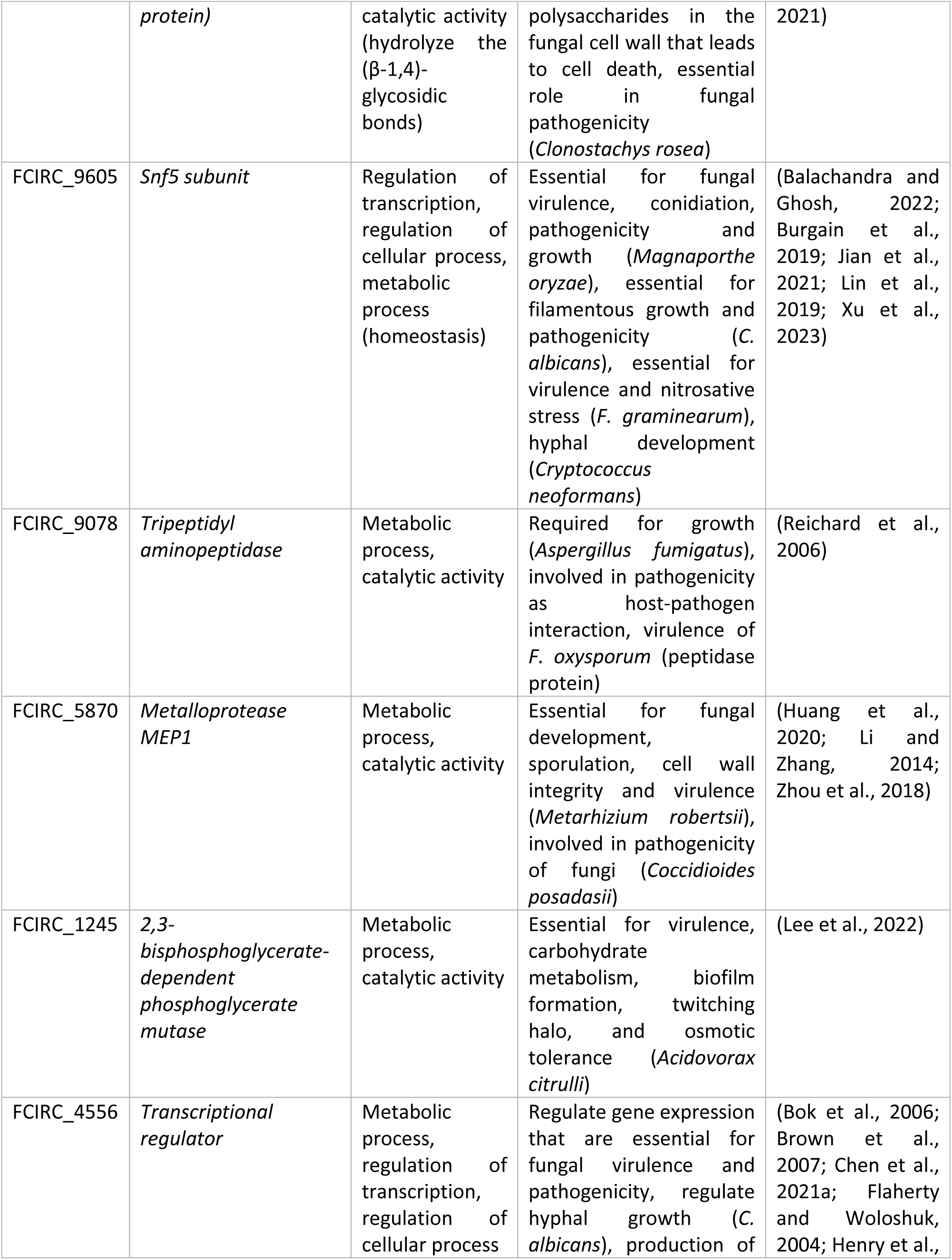
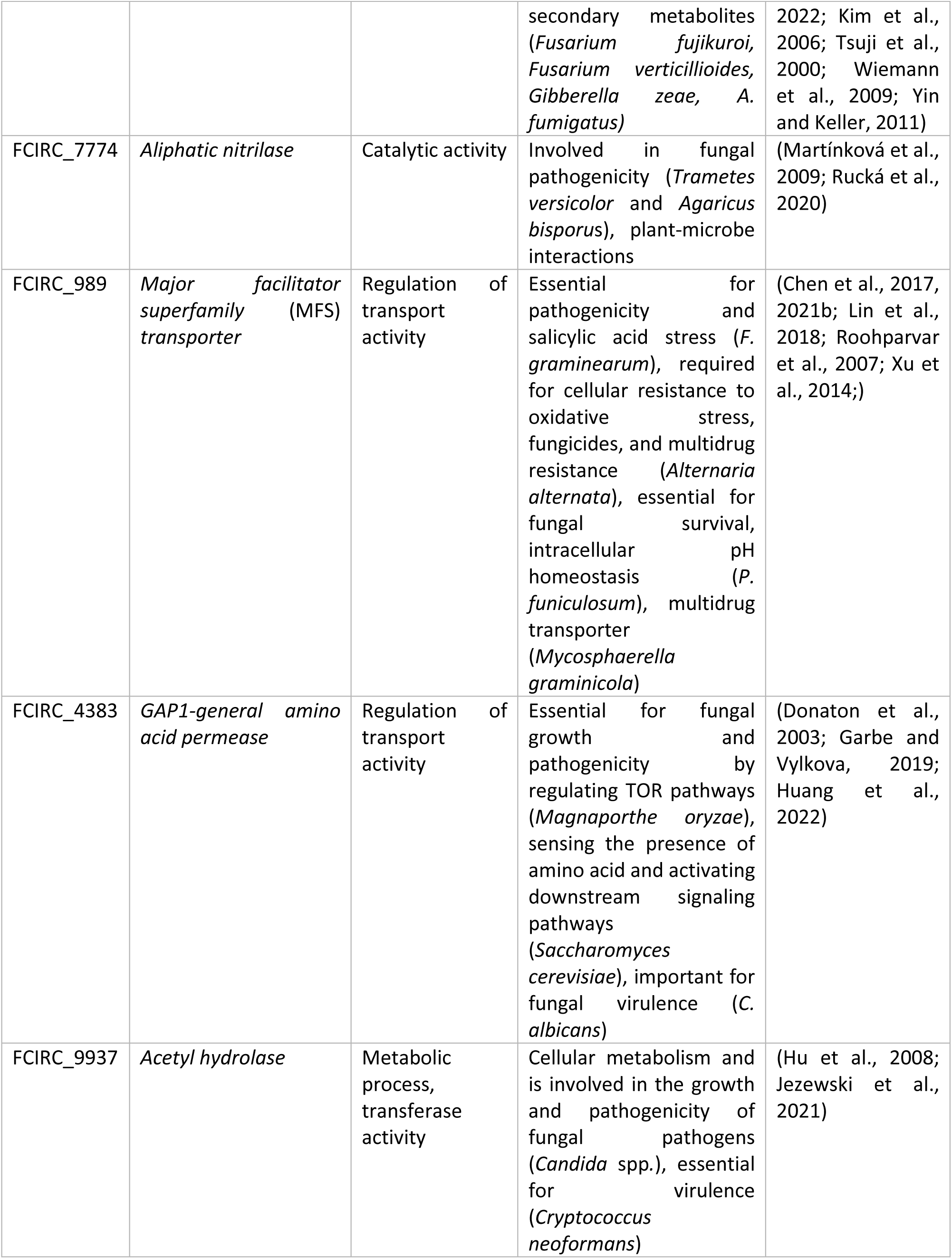
Putative Pathogenicity and Virulence associated downregulated genes in *Fusarium circinatum* after the application of exogenous dsRNA-*PTP*.

The 10 others down-regulated genes, *Exoglucanase type* (FCIRC_9483)*, GPI transamidase component PIG-S* (FCIRC_4987)*, Hypothetical protein (MutS family DNA mismatch repair protein)* (FCIRC_13497), *Positive regulator of purine utilization* (FCIRC_9481)*, Phenylacetone monooxygenase* (FCIRC_13714)*, Regulatory P domain-containing protein* (FCIRC_1034)*, β-mannosidase* (FCIRC_8458)*, Hypothetical Protein* (*TauD domain-containing protein/CAS_like super family*) (FCIRC_471)*, Eliciting plant response* (FCIRC_12765), and *Hypothetical Protein* (FCIRC_1906) and the 3 up-regulated genes, *Peptide transporter* (FCIRC_9294)*, Nuclease PA3* (FCIRC_4281), and *Acetoacetate decarboxylase* (FCIRC_1460), are essential as for conidiation, growth, host-pathogen interaction, nutrient acquisition, and genome stability of the pathogen (table S6 and table S7). Consequently, using RNA-seq promising candidate target genes to be assessed in future studies were identified.

## Discussion

Spray Induced Gene Silencing (SIGS) is a potentially sustainable and environmentally friendly disease control strategy whose success has been demonstrated in the control of several crop pathogens. In particular, SIGS can provide control against various *Fusarium* species, such as *F. graminearum* (Koch et al., 2016), *F. culmorum* (Tretiakova et al., 2022), and *Fusarium oxysporum* f. sp. *lycopersici* (Ouyang et al., 2023). Despite the many advantages of SIGS for the control of forest diseases (Bocos-Asenjo et al., 2022), it has not yet been investigated for this purpose. In this study, we aimed to determine whether exogenous application of specific dsRNAs could be an effective protection strategy against forest pathogens, taking the example of *F. circinatum*, the causal agent of PPC disease. This is thus the first report of SIGS for the control of a forest pathogen.

SIGS control of pathogens is dependent on their dsRNA uptake ability, which varies greatly between different fungal species (Qiao et al., 2021). The results of our study demonstrate that *F. circinatum* is able to efficiently uptake exogenous dsRNA to trigger RNAi. This is in accordance with previous reports with various *Fusarium* species, including *F. graminearum* (Koch et al., 2016; Qiao et al., 2021) and *F. oxysporum* f. sp. *lycopersici* (Ouyang et al., 2023).

RNAi-mediated plant protection is not only dependent on silencing efficiency, but also on the selection of appropriate target genes. Studies using RNAi for pathogen and pest control target virulence-determining or essential genes, resulting in reduced virulence as a product of functional disruption (Sang and Kim, 2020). However, many of the genes involved in virulence, such as effector genes, are expressed only for a limited period (sometimes late in the course of infection progression). Hence, applying SIGS treatments at the correct time to silence virulence genes might be challenging. In contrast, essential genes are always expressed, which makes their silencing more effective. Indeed, numerous studies have demonstrated that the knockdown through RNAi of essential genes involved in the regulation of physiological processes, including cell transport, cell division, biosynthesis of structures, etc., can lead to increased mortality of the target organism (Castellanos et al., 2018; Hernández-Soto and Chacón-Cerdas, 2021; Panwar et al., 2018). Therefore, a factor contributing to the success of this technology is the essential nature of the targeted gene (Ray et al., 2022). This is why we designed dsRNA molecules to target genes in three essential pathways for fungal development (vesicle trafficking, transduction of the signal, and cell wall biogenesis) since we found that blocking these critical pathways reduced fungal virulence and infection. This agrees with several studies that have shown that targeting essential genes in *Fusarium* spp. using RNAi strategies confers protection to the plants (Cheng et al., 2015; Chen et al., 2016a; Ghag et al., 2014; Koch et al., 2013; Machado et al., 2018; Yang et al., 2021).

Silencing multiple genes simultaneously increases the chances of effectively silencing the pathway (Koch et al., 2019; Mumbanza et al., 2013; Qiao et al., 2021). For instance, in *F. graminearum* spraying barley leaves with a dsRNA construct targeting three genes (FgCYP51A, FgCYP51B, and FgCYP51C) of the pathogen resulted in a greater reduction of the symptoms than applying single (CYP-A, CYP-B, CYP-C) and double (CYP-AC, CYP-BC, CYP-AB) dsRNA constructs (Koch et al., 2019). Besides, targeting multiple genes can decrease the likelihood of resistance or adaptation of the fungus to SIGS, as various genes may possess varying degrees of duplication or compensation in the fungal genome. Also, certain genes may possess counterparts or duplicates that can carry out comparable tasks or assume the function of the suppressed gene. Therefore, silencing such genes may not hinder the fungus from overcoming the RNAi (Padilla-Roji et al., 2023; Ray et al., 2022). Consequently, we designed dsRNA molecules that targeted more than one gene within the same pathway.

dsRNA-*VDS* targeted three genes in the vesicle trafficking pathway (*VPS51*, *DCTN1*, and *Sac1*). siRNAs targeting these genes were previously found within extracellular vesicles released by *Arabidopsis thaliana* plant cells during the infection of *B. cinerea,* leading to pathogen virulence inhibition (Cai et al., 2018). Additionally, they have been previously tested by (Qiao et al., 2021), who silenced them in several pathogenic fungi by SIGS and achieved a reduction of infection symptoms in a pathogen uptake-dependent process. In this work, we have further confirmed that these genes make useful targets for SIGS. dsRNA-*PTP* targeted four genes in the rapamycin (TOR) signaling pathway (*Pp2a*, *Sit4*, *Ppg1*, and *Tap42*). This pathway plays a critical role in nutrient signal transduction and therefore in cell growth and proliferation in filamentous fungi (Yang et al., 2022; Yu et al., 2014) and had never been disrupted by RNAi before. Rapamycin acts in the TOR signaling pathway preventing the association between the *Tap42* complex and the TOR complex while *Tap42* interacts with the type 2A phosphatases *Pp2a*, *Sit4*, and *Ppg1* (Yan et al., 2012; Yu et al., 2014). In *Fusarium graminearum*, *Pp2a* is essential for pathogen survival; *Sit4* and *Ppg1* are important for cell integrity and *Tap42* may be crucial for the fungus since no deletion mutants of this gene could be obtained (Yu et al., 2014). Some *Fusarium* species are strongly inhibited by rapamycin treatment (López-Berges et al., 2010; Teichert et al., 2006; Yu et al., 2014). Therefore, it seemed reasonable to assume using SIGS to target these genes would provide some level of protection against *F. circinatum*, as was indeed observed in this study. Finally, four genes in cell wall biogenesis were targeted by dsRNA-*CHS*: three chitin synthases (*Chs1*, Chs2 and *Chs3b*) and a glucan synthase (*Gls1*). In fungal cell walls, chitin is a major component, and disruption of chitin synthesis results in malformed and osmotically unstable fungal cells (Bowman and Free, 2006). Since chitin synthases are responsible for chitin synthesis, genes that encode for these enzymes are suitable candidates for SIGS. *Chs1* and *Chs2* have been silenced in other *Fusarium* species affecting mycelial growth and virulence (Cheng et al., 2015; Xu et al., 2010). *Chs3b* deletion appears to be lethal in *F. graminearum* and its silencing by HIGS resulted in reduced and sparse mycelial growth (Cheng et al., 2015; Liu et al., 2016). Glucan is another fundamental component of the fungal cell wall, with the central core of the cell wall consisting of a branched β-1,3-glucan cross-linked to chitin (Latgé, 2007). The production of β-glucans involves a glycosyltransferase enzyme known as β-1,3-glucan synthase which is the pharmacological target of numerous antifungal drugs. In this family of enzymes, glucan synthase 1 (*Gls1*) was found to be lethal in *Colletotrichum graminicola* and its silencing by RNAi in transgenic fungal strains produced developmental abnormalities and non-pathogenicity (Oliveira-Garcia and Deising, 2013). Chen et al., (2016a) produced HIGS transgenic plants targeting *Gls1* and infected them with *F. culmorum*, finding similar results in fungal morphology and disease reduction. Perhaps unsurprisingly, we found that *Chs1*, *Chs2*, *Chs3b* and *Gls1* made good targets for SIGS against *F. circinatum*.

The study on *F. circinatum* infection was carried out using the most susceptible species to PPC, *P. radiata.* Trials on pine seedlings are time-consuming, since the seedlings used are between 4 and 6 months old, and observation of symptoms can take up to a month. To expedite the testing process, a rapid method was developed to test dsRNAs before host testing. The infection of detached spinach leaves allowed us to test the efficacy of the designed dsRNA molecules easily and within 4 days. This approach spared us the testing of all the molecules in long-lasting assays on pine seedlings. As a result, the most effective molecules in terms of virulence reduction were used in subsequent assays in pines while the rest were discarded. Rapid tests were a fundamental tool in this study and are strongly advised for research involving forest species.

The data obtained in this work suggests that SIGS is an efficient method to reduce *F*. *circinatum* virulence and disease symptoms in pine seedlings. The designed molecules reduced the virulence of the pathogenic organism by almost 50% in spinach detached leaves, based on the average lesion size caused by *F. circinatum*. These findings are consistent with the decrease in fungal biomass in dsRNA-treated leaves. In the main host, *P. radiata*, we achieved a reduction of *F. circinatum* virulence of approximately 30% according to a disease score scale modified from Correll et al. (1991). These percentage reductions are expected when using SIGS, since these reductions are never 100%, as has been proven in numerous pathogens (Niu et al., 2021) and in other *Fusarium* species (Koch et al., 2016; Ouyang et al., 2023; Tretiakova et al., 2022).

dsRNA-*CHS* molecule, which targets the enzymes responsible for the synthesis of chitin and glucan, provided the best results in protecting pine trees. The cell wall plays a fundamental role in pathogenic fungal cells, as this structure responds to the complex interactions between pathogens and their host plants (Geoghegan et al., 2017). During the initial stages of infection, chitin is essential in the formation of infection structures (Fernandes et al., 2016). Thus, this dsRNA molecule may be preventing spore germination at an early stage, i.e. reducing the effectiveness of the primary inoculum, whereas the other dsRNAs may be slowing the fungal growth. Saito et al., (2022), studied the effect of SIGS treatment targeting chitin synthase genes on the formation of infection structures of the pathogen *Phakopsora pachyrhizi*, and they found that the dsRNA suppressed the formation of these structures in abaxial and adaxial surfaces of soybean leaves. Yang et al., (2021) also targeted genes involved in cell wall biogenesis and

achieved significant levels of *F. graminearum* growth reduction in vitro, in detached leaves and spikelets of wheat plants, and observed a decrease in toxin production by the pathogen.

In this study, dsRNAs of different lengths (492 bp and 800 bp) were successfully used to reduce pathogen virulence. There are differing findings regarding the length of dsRNA molecules: Degnan et al., (2022) and Höfle et al., (2020) obtained reduced silencing efficiencies with longer dsRNAs, whereas Werner et al., (2020) obtained higher silencing efficiencies with longer dsRNAs. *F. circinatum* can uptake dsRNAs of up to 800 bp as we observed an effect of molecules of this size in reducing the virulence.

The combined application of several molecules silencing different metabolic pathways (dsRNA-mix) did not offer higher efficiency than the application of each treatment separately. This might be because the final concentration of each molecule was only a third of that used when applied individually. This allowed to not only obtain comparable results but also avoid saturating the RNAi machinery (Yang et al., 2000). Yang et al., (2021) also observed no improvement when spraying mixtures of two or three dsRNA constructs on wheat leaves compared to the protection provided by each construct individually.

Currently, SIGS is mainly a preventive technology. In this work, we demonstrate how it can reduce PPC disease in pine seedlings. SIGS is an easily applicable technology in seedlings and would therefore be very useful in nurseries, where one of the major problems is the emergence and rapid dissemination of diseases, and in the case of PPC, a major source of spread to forests.

This study shows promising results for SIGS. However, the stability and duration of the silencing are limited due to RNA instability/degradation under environmental conditions (Hoang et al., 2022; Niño-Sánchez et al., 2022; Qiao et al., 2023). Re-applying dsRNAs regularly would be feasible in the field and might help to overcome the issues related to the loss of efficacy over time. We tested this hypothesis on pine seedlings and obtained promising results. To the best of our knowledge, this is the first study to test the repeated application of dsRNAs. Although we demonstrated that recurring dsRNAs applications improve the efficiency and durability of SIGS, we did not achieve complete disease protection or control. The use of various types of nanocarriers such as artificial vesicles (Qiao et al., 2023), clay nanoparticles (BioClay^TM^) (Mitter et al., 2017; Niño-Sánchez et al., 2022), or carbon-based (Wang et al., 2023) among others, can enhance RNA stability and uptake efficiency by the pathogens. Furthermore, it has also been shown to increase the protection window against fungal pathogens, viruses and insects (Jain et al., 2022; Mitter et al., 2017; Mosa and Youssef, 2021). Therefore, we believe that encapsulating our dsRNAs would increase their efficacy and provide prolonged protection against *F. circinatum*, and further work in that direction is needed.

Transcriptomic alterations in the *F. circinatum* transcript after applying signal-transduction dsRNA (dsRNA-*PTP*) via SIGS were determined. This research is the first to showcase the effects of targeting signal-transduction genes through SIGS, which modulates transcriptional responses. The application of dsRNA-*PTP* to silence *F. circinatum* signaling genes *Pp2a*, *Sit4*, *Ppg1,* and *Tap42* (regulating TOR Pathway) resulted in the modulation of the expression of 24 other genes that are necessary for pathogenicity and full virulence. Similarly, genes involved in signaling transduction in the TOR pathways play an important role in regulating the expression of numerous other pathways in various filamentous fungi, including hyphal growth and morphogenesis, nutrient sensing, vegetative development, cell cycle regulation, and integrity, secondary metabolism, and stress response (Fitzgibbon et al., 2005; Hou et al., 2002; Weisman et al., 2001; Yu et al., 2014). Based on the RNA-seq findings, we identified various potential targets for SIGS against *F. circinatum* that should be assessed in future studies. These essential genes might even be useful to control other fungal pathogens.

SIGS represents a cutting-edge approach with clear advantages against forest pathogens, although it has been scarcely explored so far. One of its greatest strengths is its easy spray application which allows for broad and efficient dissemination of dsRNA molecules across large forest areas. This scalable solution can combat diverse pathogens, making it an ideal solution for forest management since dsRNA production is becoming more and more economically competitive (Bocos-Asenjo et al., 2022). The specificity of this technique is also a great advantage in forests because it allows the selective silencing of pathogen genes while leaving non-target organisms undisturbed. This approach minimizes ecological alterations, preserving the delicate balance of forest ecosystems while protecting them from diseases, contrary to fungicides that can cause ecological risks to the natural environment such as contamination of groundwater, and affect a broad range of organisms. Nevertheless, using SIGS to protect forests is not without challenges due to the vast area to be covered and the large quantities of dsRNA required to ensure complete protection.

In conclusion, in this work, it is shown that application of SIGS is a promising alternative for the control of Pine Pitch Canker disease caused by *Fusarium circinatum*. The present study is a significant step toward environmentally friendly forest management and shows the technological potential of novel RNA-based biofungicides.

## Acknowledgments

This research was funded by the Spanish Ministry of Science and Innovation and by the European Union through the Next Generation funds (project numbers: PID2019-110459RB-I00; PLEC2021-008076). This study has been also funded by the Junta de Castilla y León through the projects “VAP208P20”, “VA178P23”, “CLU-2019-01 – iuFOR Unit of Excellence” of the University of Valladolid and co-funded by European Regional Development Fund (ERDF “Europe drives our growth”). J.N.S. received support from the European Union’s Horizon Europe research and innovation programme under the MSCA agreement No 101068728. I.T.B.A. and H.A. were recipients of PhD fellowships funded by Junta de Castilla y León (Orden EDU/601/2020 and Orden EDU/1508/2020). S.D.H. received a Juan de la Cierva postdoctoral fellowship funded by MCIU/AEI/10.13039/501100011033 and “NextGenerationEU”/PRTR (Spain).

