## Supplementary materials for "Spray-induced gene silencing (SIGS) as a tool for the management of Pine Pitch Canker forest disease"

### Supporting material

Table S1. Sequences of dsRNAs and plasmids used in this research.

| dsRNA sequence | dsRNA sequence Plasmid map |
|---|---|
| **GGTACC**CGGCCTGCAGTGCTTCGCCCGCTACC<br>CCGACCACATGAAGCAGCACGACTTCTTCAAG<br>TCCGCCATGCCCGAAGGCTACGTCCAGGAGCG<br>CACCATCTTCTTCAAGGACGACGGCAACTACAA<br>GACCCGCGCCGAGGTGAAGTTCGAGGGCGAC<br>ACCCTGGTGAACCGCATCGAGCTGAAGGGCAT<br>CGACTTCAAGGAGGACGGCAACATCCTGGGGC<br>ACAAGCTGGAGTACAACTACAACAGCCACAAC<br>GTCTATATCATGGCCGACAAGCAGAAGAACGG<br>CATCAAGGTGAACTTCAAGATCCGCCACAACA<br>TCGAGGACGGCAGCGTGCAGCTCGCCGACCAC<br>TACCAGCAGAACACCCCCATCGGCGACGGCCC<br>CGTGCTGCTGCCCGACAACCACTACCTGAGCTA<br>CCAGTCCGCCCTGAGCAAAGACCCCAACGAGA<br>AGCGCGATCACATGGTCCTGCTGGAGTTCGTG<br>ACCGCCGCCGGGATCACTCTCGGCATGGACGA<br>GCTGTACAAG**AGATCT** | 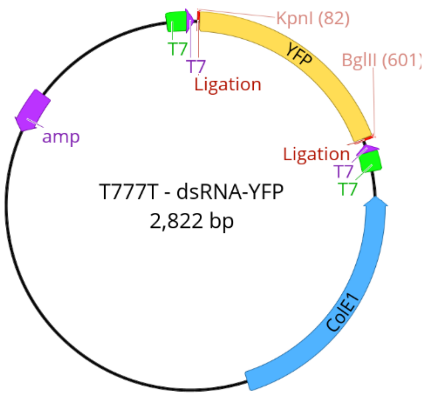 |
| **GGTACC**TCTTTCAAAGAGCTTGAAGATCAAATC<br>CAGATTCTCTGGGATGAACGGGCTGAGTCATC<br>GCACAACGGTAAAGTGGCCATGTACTTCCTCC<br>GTGTTTTACGAGATATCCGACATCAGTTACCAG<br>AGCGCCCTTCCATCAAGAGCTTTGGCCTGAAG<br>GCTGTTCCTTCACAATCTCTTCTTGAAAAGAAG<br>AACAGTCGCGAGGACGAGCTCGAGAAGGAGC<br>TTGAAACAATCCGTCAGAACGGTACAGCGGCC<br>AACAGGTCTTCTTTCAGAGACTCCCGAGACACC<br>GTCGTAATGGCCCAGGCAATGGAGAACCGACC<br>AGTCGAGAACCTCAAGGTCTGCCAGAGCTCAT<br>TCGCAAAGCATATGCTGGAAGTTCAACTCAAG<br>GAGGAAGGCTTTGACATGAGTGCGCAGTCTGA<br>CCAAGTGACTGCTTGGTTCAACACCCTCTGGGC<br>GGACAATGGCGACGCTGTGTCAAAACAGTACG<br>CATCGACAGCAGCGATGAAG**AGATCT** | 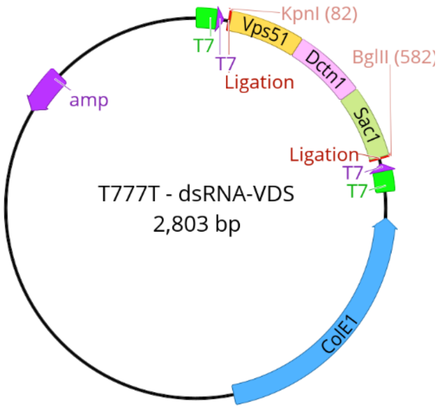 |

<table>
<tr>
<td>GGTACCACCCTCAACGAATCACAATCCTCCGA<br>GGAAACCACGAGTCCCGTCAGATCACTCAGGT<br>CTACGGTTTCTACGACGAATGCCTCCGTAAATA<br>TGGAAACGCTAATGTCTGGAAGTATTTTACCG<br>ACCTCTTTGACTACCTCCCCCTCACTGCCCTGAT<br>CGACAACCAAATCTTCTGTCTTCATGGCGGCCT<br>CTCCCCAAGCATGTTCGACTTTTTAACTTTGAG<br>CGTTGTGATCAACGACCAAATATTTTGCGTCCA<br>TGGAGGCCTTTCCCCTTCTATCCATTCTATCGAC<br>CAGATCAAAATTATAGATCGTTTCCGCGAGATC<br>CCCCATGAAGGTCCCATGGCCGATCTCGTCTG<br>GTCTGATCCTGACCCTGAGCGCGACGAATTCTC<br>CCTTTCACCTCGCCAACAGAGATCACAAACCCC<br>AAGTTGAGGAAGAAGATAAAGTCTTCGGGCG<br>AGAATGCAACATCCAGCGGCGTGGAGGAGAC<br>CGAGGCAGTCGAGGCAGAGGAAGATCCTGAT<br>TCCACAATGGCAGACCAGACCGAATCTGGCGT<br>CACAGTCAACAAGTCAAGTCAAAGTGCCGATA<br>CTCGATACATTTTCCTGGGAGATATCCCTCTAC<br>TCTTCTGCGCGCGACCAAGTCTCAGCTCTTGCG<br>CTCTTTTCCTCCAACGAAGGACTTGAGGACGTT<br>TCTACATCAGATCTACCTTATCTGCTTCTCGACG<br>CCCACGTTGCGGAGCTCGTACAAAAGACCCCC<br>AATCAAAGCCCTGACCAACGCCTAGAGGTCCT<br>CTCCAAGTCTCGCGCCGCCTACGAGCCGCGG</td>
<td>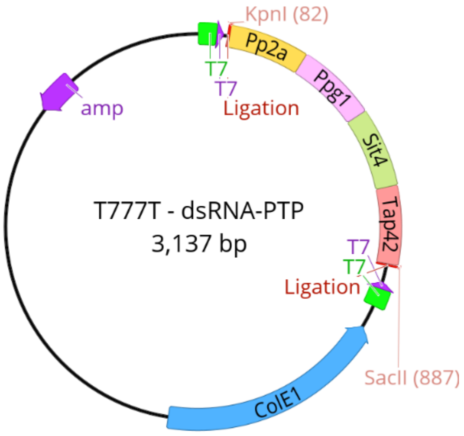<br></td>
</tr>
<tr>
<td>GGTACCCGTCACATTCTCCTGGTTCTCACTCGC<br>GTCTTTCTACCTCACCACCACCATCATCATGAA<br>GCTTGTCGGAACGCCCCAGGTTCTCTCCGGAT<br>ACCATGGTTGGCCATTTGGCGATACAGCCACG<br>CCGATTGTTAATGTCCTGATCAAATATATATAC<br>ATCGCCTTCCTCGTCCTCCAATTCGTCCTCGCTC<br>TCGGAAACACGAAGAACACTACGACCCTCATA<br>CGTATGTACCACCCTCATCACCCAGCCAGATTC<br>AGACACAGACACCTCCTGCACGCCCTCCCTCCT<br>CTCCTTCTGTTTCCCCTTCTACCTCATGTTCAAA<br>GACAACACCCTCCCCTCTCAAACCGCCAGTAGC<br>GTTTCCTCCTCCATTGTCAATTTCACCTGTGAGC<br>ATCACCATGTTTCCTCGACTTCCTCCGATCCGAT<br>CACACTTTCTTGCGAAAGTTTGCCTTCTTCATCG<br>AATTCATCTTCAACACTATCAACATGATCTTCGC<br>TTGGTTTGCTATTGGTAACTTCTTCCTCGTTTTC<br>AAGATTCTCACAACAAGTTTGGGAGACGATTC<br>GTTACTGGGGCGGACAGGAGAAATTCTAGGT<br>GTCGTATTTACACTACCAAGATGATCAGTACTA<br>CGATCAAGGCTACGACAACCGTGGCCCAAATC<br>ACAATAACAACAATAACAACAATAACCATGAT<br>GGCTACTACGATGAATCTGGTTACTACAACGCC<br>GACCCAAATAACCCCTACCAGCAGGATGGAGG</td>
<td>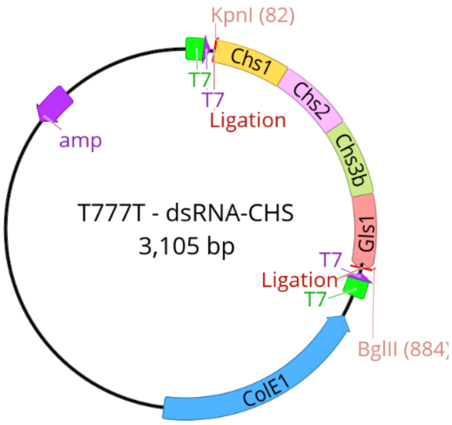<br></td>
</tr>
</table>

| |
|---|
| CTATTACGACGGCCACGATCAATACCAAGATG GATACTACGACAATGGCAGATCT |

Sequences underlined with different colors represent fragments of various genes that were used as dsRNA templates. Letters in red correspond to the restriction enzymes.

Table S2. qPCR primers used for Fungal Biomass Quantification

| **Primer name** | **Primer sequence (5'-3')** |
|---|---|
| *FcRPB2*-q-F | ACCCGTCTTCACAGTTCACC |
| *FcRPB2*-q-R | TGTCTTCAATCGCTTGTTGC |
| *SoActin*-q-F | TGTTGCCCCAGAAGAGCACCCT |
| *SoActin*-q-R | AGAACGGCCTGAATGGCAACATAC |

Table S3. Statistical summary corresponding to figure 4A.

| **Centrality and dispersion estimations figure 4A** | | | | | |
|---|---|---|---|---|---|
| **Treatment** | **Dpi** | **Mean** | **SD** | **Median** | **IQR** |
| Water | 25 | 0 | 0 | 0 | 0 |
| *F. circinatum* | 25 | 2.39 | 0.49 | 2.5 | 0.75 |
| dsRNA-*YFP* | 25 | 2.54 | 0.57 | 2.5 | 1 |
| dsRNA-*CHS* | 25 | 1.49 | 0.77 | 1.5 | 1 |
| dsRNA-*PTP* | 25 | 1.67 | 0.80 | 2 | 1 |
| dsRNA-*VDS* | 25 | 2.01 | 0.84 | 2 | 1 |
| dsRNA-mix | 25 | 1.80 | 0.80 | 2 | 1.5 |
| Water | 35 | 0.00 | 0.00 | 0 | 0 |
| *F. circinatum* | 35 | 3.74 | 0.67 | 4 | 0.75 |
| dsRNA-*YFP* | 35 | 3.77 | 0.61 | 4 | 0.5 |
| dsRNA-*CHS* | 35 | 2.90 | 0.57 | 3 | 0.75 |
| dsRNA-*PTP* | 35 | 3.07 | 0.73 | 3 | 0.5 |
| dsRNA-*VDS* | 35 | 3.21 | 0.85 | 3.5 | 1.375 |
| dsRNA-mix | 35 | 3.24 | 0.72 | 3.5 | 0.5 |
| **Statistical analysis figure 4A** | | | | | |
| **Comparison** | **N** | **Mean diff** | **lwr.ci** | **upr.ci** | ***p*-value** |
| 25 dpi *Fcir* vs *YFP* | 35 vs 35 | 0.16 | -0.28 | 0.60 | 0.83 |
| 25 dpi *Fcir* vs *CHS* | 35 vs 35 | -0.90 | -1.34 | -0.46 | 2e-6 |
| 25 dpi *Fcir* vs *PTP* | 35 vs 35 | -0.71 | -1.15 | -0.28 | 0.0002 |

| | | | | | |
|---|---|---|---|---|---|
| 25 dpi *Fcir* vs *VDS* | 35 vs 35 | -0.37 | -0.81 | 0.07 | 0.013 |
| 25 dpi *Fcir* vs mix | 35 vs 35 | -0.59 | -1.02 | -0.15 | 0.004 |
| 25 dpi *YFP* vs *CHS* | 35 vs 35 | -1.06 | -1.50 | -0.62 | 2e-8 |
| 25 dpi *YFP* vs *PTP* | 35 vs 35 | -0.87 | -1.31 | -0.43 | 4.8e-6 |
| 25 dpi *YFP* vs *VDS* | 35 vs 35 | -0.53 | -0.97 | -0.09 | 0.013 |
| 25 dpi *YFP* vs mix | 35 vs 35 | -0.74 | -1.18 | -0.30 | 0.0001 |
| 35 dpi *Fcir* vs *YFP* | 35 vs 35 | 0.03 | -0.39 | 0.45 | 0.99 |
| 35 dpi *Fcir* vs *CHS* | 35 vs 35 | -0.84 | -1.26 | -0.42 | 3.9e-6 |
| 35 dpi *Fcir* vs *PTP* | 35 vs 35 | -0.67 | -1.09 | -0.25 | 0.0003 |
| 35 dpi *Fcir* vs *VDS* | 35 vs 35 | -0.54 | -0.96 | -0.11 | 0.007 |
| 35 dpi *Fcir* vs mix | 35 vs 35 | -0.50 | -0.92 | -0.08 | 0.013 |
| 35 dpi *YFP* vs *CHS* | 35 vs 35 | -0.87 | -1.29 | -0.45 | 2.5e-6 |
| 35 dpi *YFP* vs *PTP* | 35 vs 35 | -0.70 | -1.12 | -0.28 | 0.0001 |
| 35 dpi *YFP* vs *VDS* | 35 vs 35 | -0.57 | -0.99 | -0.14 | 0.004 |
| 35 dpi *YFP* vs mix | 35 vs 35 | -0.53 | -0.95 | -0.11 | 0.007 |

Table S4. Statistical summary corresponding to figure 5A.

| **Centrality and dispersion estimations figure 5A** | | | | | |
|---|---|---|---|---|---|
| **Treatment** | **Type** | **Mean** | **SD** | **Median** | **IQR** |
| *F. circinatum* | weekly | 3.20 | 0.79 | 3.50 | 1.38 |
| dsRNA-*YFP* | weekly | 3.55 | 0.80 | 3.50 | 0.88 |
| dsRNA-*CHS* | weekly | 2.23 | 0.82 | 2.50 | 0.50 |
| **Statistical analysis figure 5A** | | | | | |
| **Comparison** | **N** | **Mean diff** | **lwr.ci** | **upr.ci** | ***p*-value** |
| Weekly *YFP* vs Control | 30 vs 30 | 0.35 | -0.11 | 0.81 | 0.16 |
| Weekly *CHS* vs Control | 30 vs 30 | -0.96 | -1.43 | -0.49 | 2.3e-5 |
| Weekly *YFP* vs *CHS* | 30 vs 30 | -1.31 | -1.78 | -0.84 | 2e-8 |

Table S5. Statistical summary corresponding to figure S2

| **Centrality and dispersion estimations figure S2** | | | | | |
|---|---|---|---|---|---|
| **Treatment** | **Type** | **Mean** | **SD** | **Median** | **IQR** |
| dsRNA-*CHS* | single | 2.83 | 0.95 | 2.5 | 1.38 |
| dsRNA-*CHS* | weekly | 2.23 | 082 | 2.5 | 0.5 |
| **Statistical analysis figure S2** | | | | | |
| **Comparison** | **N** | **Mean diff** | **lwr.ci** | **upr.ci** | ***p*-value** |
| *CHS* single vs Weekly | 30 vs 30 | -0.6 | -1.13 | -0.067 | 0.025 |

Table S6. Essential Downregulated genes/proteins in *Fusarium circinatum* after the application of exogenous dsRNA-*PTP.*

| **Accession No.** | **Putative Gene/Proteins** | **Function** | **Descriptive characteristics** | **References** |
|---|---|---|---|---|
| FCIRC_9483 | *Exoglucanase type C* | Catalytic activity (hydrolysis of cellulose), cell development | Adhesion to host-cell surfaces (*Candida albicans*), morphogenetic-morpholytic processes during development and differentiation, promotes lactate-induced β-glucan masking (*C.albicans*), influence immune to other fungal responses | (Martin et al., 2007; Tsai et al., 2011; Childers et al., 2020; Rafiei et al., 2021) |
| FCIRC_4987 | *GPI transamidase component PIG-S* | Biosynthetic process | Crucial role in the GPI anchoring process in fungi, essential for growth (*Saccharomyces cerevisiae*) | (Ohishi et al., 2001; Kinoshita and Fujita, 2016) |
| FCIRC_13497 | *Hypothetical protein (MutS family DNA mismatch repair protein)* | Regulate cell growth, regulate genome stability | Essential for genome stability | (Chen et al., 2016b) |
| FCIRC_9481 | *Positive regulator of purine utilization* | Regulation of transcription, regulation of cellular processes | Crucial role in uptake and utilization of purine in various fungi, regulates uracil degradation *(S. cerevisiae*), purine utilization pathway (*Aspergillus nidulans*), regulation of purine utilization (*C. albicans*) | (Pantazopoulou and Diallinas, 2007; Gournas et al., 2011; Chitty and Fraser, 2017) |

| | | | | |
|---|---|---|---|---|
| FCIRC_13714 | *Phenylacetone monooxygenase* | Metabolic process, catalytic activity | Involved in sporulation of (*Colletotrichum gloeosporioides s.s. and Magnaporthe oryzae)* | (Wang et al., 2020) |
| FCIRC_1034 | *Regulatory P domain-containing protein* | Regulate cell growth, regulate gene transcription | Unknown | (Zhou et al., 1998) |
| FCIRC_8458 | *β--mannosidase* | Catalytic activity | Unknown | (Alkhayat et al., 1998) |
| FCIRC_471 | *Hypothetical protein (TauD domain-containing protein/CAS_like super family)* | Metabolic process (Oxidoreductase) | Unknown | (Eichhorn et al., 1997) |
| FCIRC_12765 | *Eliciting plant response* | Unknown function | Unknown | |
| FCIRC_1906 | *Hypothetical Protein* | Unknown function | Unknown | |

Table S7. Essential Upregulated genes/proteins in *Fusarium circinatum* after the application of exogenous dsRNA-*PTP*.

| **Accession No.** | **Putative Gene/Proteins** | **Function** | **Descriptive characteristics** | **References** |
|---|---|---|---|---|
| FCIRC_9294 | *Peptide transporter* | Regulation of transport of peptides | Nutrient acquisition in fungi (*Candida albicans*), required for growth (*Candida albicans*) | (Dunkel et al., 2013) |
| FCIRC_4281 | *Nuclease PA3* | Catalytic activity | Unknown | |
| FCIRC_1460 | *Acetoacetate decarboxylase* | Metabolic process, catalytic activity | Unknown | |

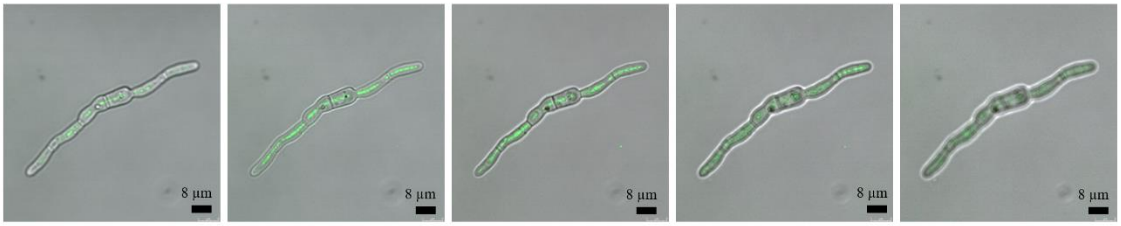

Figure S1. Z stack confocal images of a germinated spore of *Fusarium circinatum* treated with fluorescein labeled dsRNA showing the presence of fluorescence inside the germling, demonstrating that the pathogen is able to uptake external dsRNA molecules. The complete Z stack consisted of 5 images across a depth of 8.64 µm. Images were taken on a Leica Microsystems SP8 system; scale bars correspond to 8 µm.

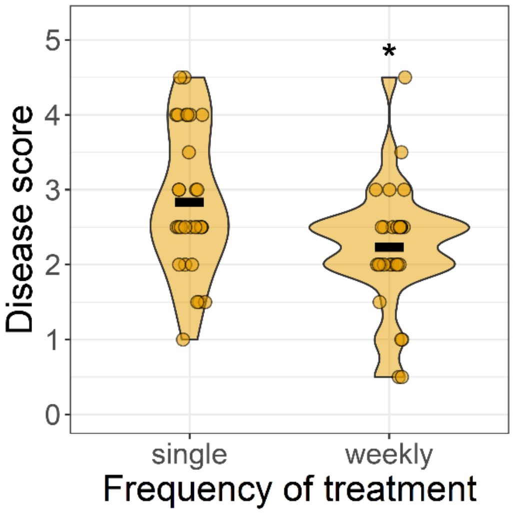

Figure S2. Sequential treatment with dsRNA-*CHS* decreased *Fusarium circinatum* virulence in pine seedlings. *Pinus radiata* seedlings were inoculated with *F. circinatum* and treated with dsRNA-*CHS* on the day of inoculation and weekly, disease severity was assessed at 30 dpi according to the modified scale of Correl (1991). Data distribution was represented through violin plots, black horizontal bars depict the mean disease score, and statistically significant differences were found by OLR test and are represented by asterisks (*: $p < 0.05$).

### Data Availability Statement

Transcriptome data are available at NCBI database PRJNA1051661.
